# Hedgehog and Bmp signaling pathways play opposing roles during establishment of the cardiac inflow tract in zebrafish

**DOI:** 10.1101/2025.07.19.665705

**Authors:** Rhea-Comfort A. Robertson, Hannah G. Knight, Catherine Lipovsky, Jie Ren, Neil C. Chi, Deborah Yelon

**Affiliations:** Department of Cell and Developmental Biology, School of Biological Sciences, University of California, San Diego, La Jolla, CA, 92093, USA; Division of Cardiovascular Medicine, Department of Medicine, University of California, San Diego, La Jolla, CA, 92093, USA

**Keywords:** pacemaker cells, cardiac specification, Islet1, *smoothened*, *acvr1l*, *chordin*

## Abstract

Cardiac pacemaking activity is controlled by specialized cardiomyocytes in the cardiac inflow tract (IFT), but the processes that determine IFT dimensions remain poorly understood. Here, we show that Hedgehog (Hh) signaling limits the number of IFT cardiomyocytes in the embryonic zebrafish heart. Inhibiting Hh signaling, either genetically or pharmacologically, results in an expanded IFT population. In contrast, reducing Bmp signaling decreases the number of IFT cardiomyocytes, while increasing Bmp signaling leads to an excess of IFT cardiomyocytes. Temporal inhibition of each pathway reveals that Hh and Bmp signaling act before myocardial differentiation to regulate IFT size. Simultaneous reduction of both Hh and Bmp signaling yields a relatively normal number of IFT cardiomyocytes, suggesting that these pathways function antagonistically during IFT development. Additionally, epistasis analysis suggests that Bmp signaling acts upstream of Wnt signaling to promote IFT formation, whereas Hh signaling limits IFT size in a Wnt-independent manner. Our results support a model in which Hh signaling restricts the establishment of the IFT progenitor pool, while Bmp signaling drives IFT progenitor specification prior to Wnt-directed IFT differentiation.

**SUMMARY STATEMENT:** Hedgehog signaling acts prior to myocardial differentiation to restrict the number of inflow tract cardiomyocytes, while Bmp signaling acts during a similar timeframe to promote inflow tract cardiomyocyte formation.

## INTRODUCTION

The heart is composed of multiple types of specialized cardiomyocytes, each with distinct functions. Cardiomyocyte diversification begins with the establishment of discrete pools of cardiac progenitor cells, but the patterning processes that specify each progenitor population remain incompletely understood. For example, cardiac pacemaking activity is confined to a small population of cardiomyocytes that possess specific conductive properties and are located near the sinoatrial junction (Christoffels et al., 2010; van Weerd and Christoffels, 2016; Burkhard et al., 2017; Bhattacharyya and Munshi, 2020). Although the highly specialized nature of pacemaker cardiomyocytes is well established, we lack a full picture of the mechanisms that control the specification of pacemaker progenitor identity.

Pacemaker cardiomyocytes originate from late-differentiating cardiac progenitors that first reside at the periphery of the anterior lateral plate mesoderm (ALPM) and are later appended to the venous pole of the atrium (Mommersteeg et al., 2010; Bressan et al., 2013; Liang et al., 2013; Fukui et al., 2018; Ren et al., 2019). In this *Tbx18*-expressing population, a transcription factor network, including Tbx18, Tbx3, Isl1, and Shox2, directs pacemaker cardiomyocyte differentiation (Mommersteeg et al., 2010; Bakker et al., 2012; Tessadori et al., 2012; Liang et al., 2013; van Weerd and Christoffels, 2016; Martin and Waxman, 2021). Canonical Wnt signaling serves as a key positive regulator of the pacemaker differentiation program (Bressan et al., 2013; Ren et al., 2019; Liang et al., 2020). However, it is still unclear which signaling pathways act upstream of this program to allocate an appropriate number of cardiac progenitor cells into the pacemaker lineage.

The zebrafish is a useful model organism for investigating the regulation of pacemaker cardiomyocyte development (Tessadori et al., 2012; Burkhard et al., 2017; Martin and Waxman, 2021). In the embryonic zebrafish heart, pacemaker cells are located within the atrial inflow tract (IFT), in a region that acts as the functional equivalent of the mammalian sinoatrial node (SAN) (Arrenberg et al., 2010; Tessadori et al., 2012; Burkhard et al., 2017). (Hereafter, we refer to pacemaker cardiomyocytes as “IFT cardiomyocytes” in the zebrafish context.) Like the mammalian SAN, the zebrafish IFT initiates the heartbeat, demonstrates pacemaking activity, and expresses genes that encode key components of the pacemaker differentiation program (Arrenberg et al., 2010; Tessadori et al., 2012; Poon et al., 2016; Burkhard et al., 2017; Burkhard and Bakkers, 2018; Ren et al., 2019; Minhas et al., 2021). Furthermore, Wnt signaling is active at the lateral edges of the zebrafish ALPM, in the territory containing IFT progenitor cells, and promotes IFT cardiomyocyte differentiation in these regions (Ren et al., 2019). Thus, since many key elements of pacemaker development are highly conserved, the zebrafish provides valuable opportunities for identifying signals that guide the initial steps of pacemaker progenitor specification.

In this study, our use of the zebrafish reveals previously unappreciated roles for both Hedgehog (Hh) signaling and bone morphogenetic protein (Bmp) signaling during early steps of IFT cardiomyocyte development. Prior work has implicated both pathways in other aspects of cardiac progenitor specification. For instance, Hh signaling supports the initial formation of precardiac mesoderm. In both mouse and zebrafish, mutations in *smoothened (smo)*, which encodes the transmembrane protein responsible for Hh signal transduction, result in formation of a small heart due to early defects in mesoderm establishment (Zhang et al., 2001; Thomas et al., 2008; Guzzetta et al., 2020; Zhang et al., 2021). In mouse, Hh signaling has been shown to promote the migration of mesodermal progenitor cells during gastrulation (Guzzetta et al., 2020), and, in zebrafish, Hh is required cell-autonomously during gastrulation to maximize the number of mesodermal cells that adopt cardiac fate (Thomas et al., 2008). Hh signaling is also involved in regulating the contributions of late-differentiating progenitor cells from the second heart field (SHF). In mouse and zebrafish, Hh signaling promotes formation of the outflow tract (OFT) from SHF progenitor cells (Washington Smoak et al., 2005; Lin et al., 2006; Goddeeris et al., 2008; Hami et al., 2011), and, in mouse, Hh signaling is required for the contributions of posterior SHF-derived cells to the atrial septum (Goddeeris et al., 2008; Hoffmann et al., 2009; Xie et al., 2012; Briggs et al., 2016). These phenotypes may reflect an impact of Hh signaling on the timing of SHF progenitor cell differentiation (Rowton et al., 2022). Despite the attention paid to the effects of Hh signaling on cardiac progenitor cells in these contexts, the impact of Hh signaling on the pacemaker progenitor lineage has not been specifically elucidated.

Bmp signaling also contributes to both the early establishment of precardiac mesoderm and the later contributions of SHF progenitor cells. In species from *Drosophila* to mouse, Bmp signaling plays a role in initiating the specification of cardiac identity and the expression of precardiac mesoderm genes like *Nkx2-5* (e.g. (Frasch, 1995; Zhang and Bradley, 1996; Shi et al., 2000; Reiter et al., 2001). This involvement in mesodermal patterning is likely coupled with the broader function of Bmp signaling in the initial patterning of the embryonic dorsal-ventral axis (Langdon and Mullins, 2011; Yan and Wang, 2021). Additionally, Bmp signaling contributes to the regulation of the onset of myocardial differentiation (Yuasa et al., 2005; de Pater et al., 2012; Cai et al., 2013; Strate et al., 2015). Later, in the SHF, Bmp signaling supports the differentiation of SHF cells that contribute to the OFT (Hutson et al., 2010), as well as the proliferation of SHF cells at the venous pole that contribute to septation (Briggs et al., 2013). Prior investigations have not revealed whether Bmp signaling plays an early role in the initial specification of the pacemaker lineage. However, studies in zebrafish hint at this possibility: inhibition of Bmp signaling following gastrulation, while myocardial differentiation is underway, reduces the number of IFT cardiomyocytes (Fukui et al., 2018).

Here, we use genetic and pharmacological approaches to uncover important and early roles for Hh and Bmp signaling in defining the size of the zebrafish IFT. In contrast to other roles of the Hh pathway in promoting cardiomyocyte production, we find that Hh signaling restricts formation of IFT cardiomyocytes: reduced Hh activity results in an enlarged IFT. This phenotype reflects a requirement for Hh signaling prior to cardiac differentiation, suggesting a repressive role of the Hh pathway during IFT progenitor specification. Additionally, we show that Bmp signaling promotes IFT cardiomyocyte formation during a similar timeframe: reduced Bmp activity results in a diminished IFT, and increased Bmp signaling enlarges the IFT. Intriguingly, reducing both Hh and Bmp signaling restores the IFT to a nearly normal size, suggesting that Hh and Bmp signaling act in opposition to each other to set IFT dimensions. Finally, we show that Bmp signaling acts upstream of Wnt signaling to promote IFT cardiomyocyte production, whereas Hh signaling restricts IFT formation through a Wnt-independent pathway. Synthesizing these data, we propose that Hh and Bmp act in opposing pathways to pattern the cardiac progenitor pool and specify an appropriate proportion of IFT progenitor cells.

## RESULTS

### Loss of Hh signaling causes expansion of the cardiac IFT

While examining the hearts of *smo* mutant embryos, we were intrigued to find expanded expression of several genes that are normally found in the IFT at 48 hours post fertilization (hpf) (Fig. 1A-H). For example, *bmp4*, typically expressed in a narrow ring of IFT cardiomyocytes at the venous pole of the wild-type atrium (Fig. 1A), is expressed across a larger territory in *smo* mutants, extending from the venous pole of the atrium upward into the atrial myocardium (Fig. 1B). Similar expansions of the *hcn4, tbx18,* and *shox2* expression patterns are also evident in *smo* mutants (Fig. 1C-H).

**Figure 1.**
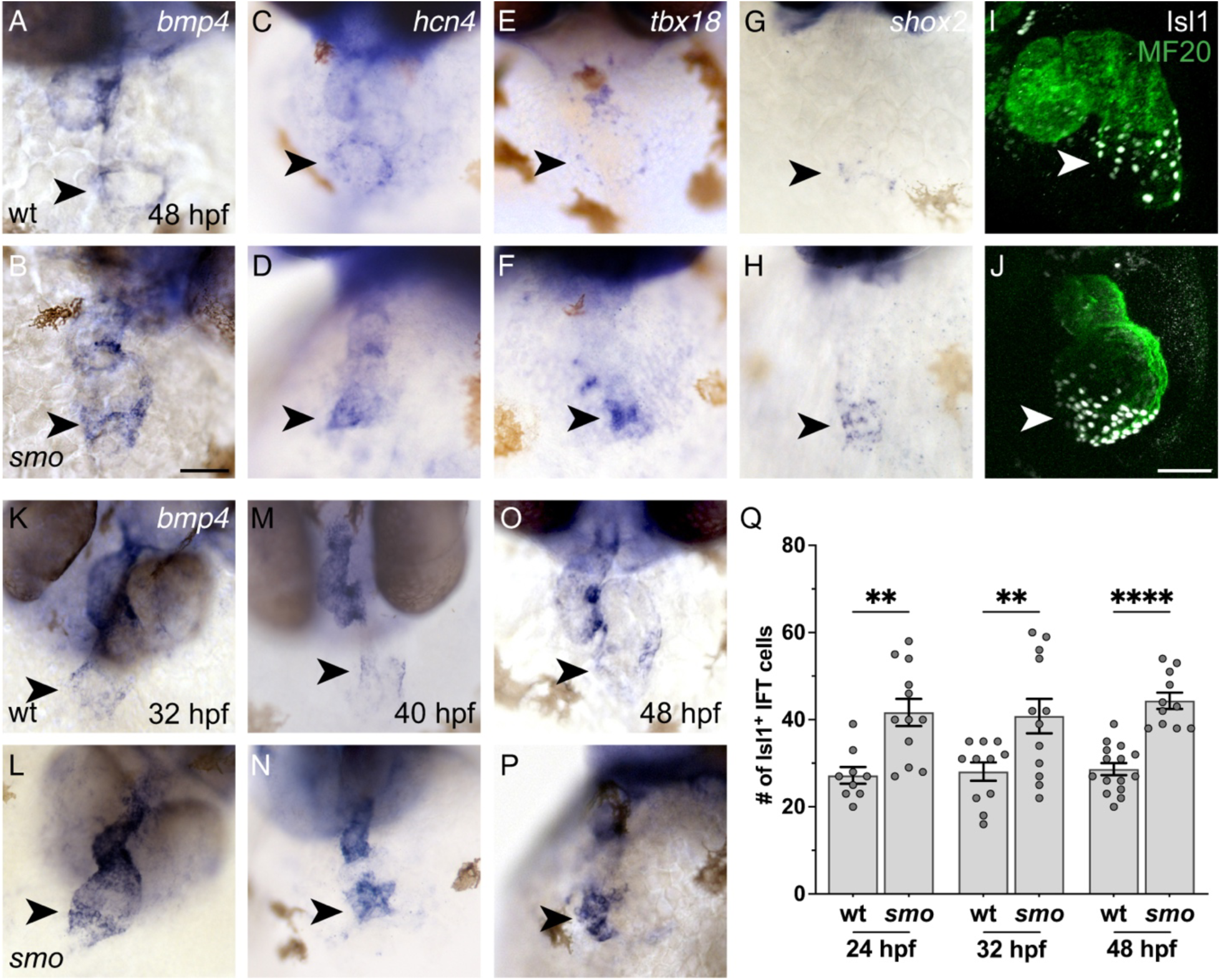
The IFT is expanded in *smo* mutants by 24 hpf. **(A-J)** Wild-type (wt; A,C,E,G,I) and *smo* mutant (B,D,F,H,J) hearts at 48 hpf are shown after in situ hybridization (A-H) or after immunofluorescence (I,J). Frontal views; arrowheads indicate the IFT. (A,B) *bmp4* expression is expanded in the venous pole of *smo* mutants relative to wt siblings (n=8 wt, 12 *smo*). (C-H) *hcn4, tbx18,* and *shox2* are similarly expanded in the venous pole of *smo* (n=10 wt, 12 *smo* for *hcn4;* n=12 wt, 12 *smo* for *tbx18*; n=26 wt, 13 *smo* for *shox2*). (I,J) Isl1 (white) is present in the nuclei of IFT cardiomyocytes; myocardium is labeled with MF20 (green). More Isl1+ cardiomyocytes are observed in the IFT of *smo* mutants than in wt siblings (n=15 wt, 11 *smo*). Scale bars: 50 μm. **(K-P)** In situ hybridization depicts *bmp4* expression in wt (K,M,O) and *smo* (L,N,P). Frontal views; arrowheads indicate the IFT. Expression of *bmp4* is already expanded in the IFT of *smo* mutants, relative to wt siblings, at 32 hpf (K,L; n=6 wt, 8 *smo*). This expansion is maintained at 40 (M,N; n=8 wt, 10 *smo*) and 48 hpf (O,P; n=8 wt, 12 *smo*). **(Q)** Graph indicates the number of Isl1+ cardiomyocytes in the IFT. See Materials and Methods for cell counting technique. The number of Isl1+ cardiomyocytes in *smo* mutants, relative to wt siblings, is increased as early as 24 hpf. This increase is maintained at 32 and 48 hpf. The number of Isl1+ cardiomyocytes in wt is also steadily maintained throughout this timeframe. **p<0.01; ****p<0.0001.

Additionally, using the transcription factor Isl1 as a marker of IFT cardiomyocytes, we found that *smo* mutants display a significant increase in Isl1+ IFT cells at 48 hpf (Fig. 1I,J,Q). This phenotype is particularly striking because the total number of cardiomyocytes is substantially smaller in *smo* mutants than it is in wild-type (Thomas et al., 2008). Thus, the expansion of the IFT cardiomyocyte population in *smo* mutants occurs in the context of a general reduction in heart size.

Since cells in the IFT act as the cardiac pacemaker, we examined heart rate and rhythm in *smo* mutant embryos. In addition to occasional arrhythmia, we found that *smo* mutants exhibit a significantly reduced heart rate compared to their wild-type siblings (Table S1). It is unclear whether these defects in *smo* mutants arise due to their expanded IFT or due to other myocardial defects. However, in mouse, failure to restrict expression of pacemaker markers can result in atrial arrhythmia (Wang et al., 2010), suggesting that a similar cause could underlie the functional defects in *smo* mutants. Altogether, these data reveal novel requirements for Hh signaling in restricting IFT size and in regulating cardiac function.

### Hh signaling limits IFT size prior to the onset of myocardial differentiation

Previous studies in zebrafish have shown that mechanisms regulated by the transcription factors Nr2f1a and Nkx2.5 act after heart tube assembly to prevent *isl1* expression from expanding beyond the venous pole of the atrium (Colombo et al., 2018; Martin et al., 2023). We therefore wondered whether the *smo* mutant phenotype might reflect a progressive expansion of Isl1+ cardiomyocytes between 24 and 48 hpf. In wild-type, the number of Isl1+ IFT cardiomyocytes remains consistent between 24 and 48 hpf (Fig. 1Q). In *smo* mutants, the number of Isl1+ IFT cardiomyocytes is increased at both 24 and 32 hpf, and the degree of IFT expansion in *smo* appears consistent through 48 hpf (Fig. 1Q). Similarly, we observed a qualitatively consistent expansion of *bmp4* expression at the IFT in *smo* mutants at 32, 40, and 48 hpf (Fig. 1K-P). These phenotypes suggest Hh signaling acts early, prior to heart tube formation, to restrict IFT dimensions.

Next, we shifted our attention to earlier stages. Our previous studies have suggested that cardiac progenitor specification occurs during gastrulation and early somitogenesis stages in zebrafish (Keegan et al., 2005; Marques et al., 2008; Thomas et al., 2008; Marques and Yelon, 2009), prior to the onset of myocardial differentiation around the 13 somite stage (Yelon et al., 1999). To test whether Hh signaling delimits the IFT population during these cardiac specification stages, we utilized cyclopamine (CyA), a potent pharmacological inhibitor of Smo (Cooper et al., 1998). Initiating CyA treatment of wild-type embryos during gastrulation (at dome stage) or at the end of gastrulation (at tailbud stage) resulted in a significant increase in the number of Isl1+ IFT cardiomyocytes at 48 hpf (Fig. 2G), similar to the increase seen in *smo* mutants (Fig. 1Q). In contrast, CyA treatment beginning at the 3 somite stage (3 s) or later did not lead to a statistically significant change in the number of Isl1+ IFT cardiomyocytes (Fig. 2G). Similarly, CyA treatment at tailbud stage resulted in expanded *bmp4* expression at the IFT (Fig. 2D,E), as seen in *smo* mutants (Fig. 1A,B,O,P), whereas CyA treatment at 10 s had no evident effect on *bmp4* expression (Fig. 2D,F). These data indicate that Hh signaling acts prior to myocardial differentiation to restrict IFT cardiomyocyte number, potentially by setting limits on IFT progenitor specification.

**Figure 2.**
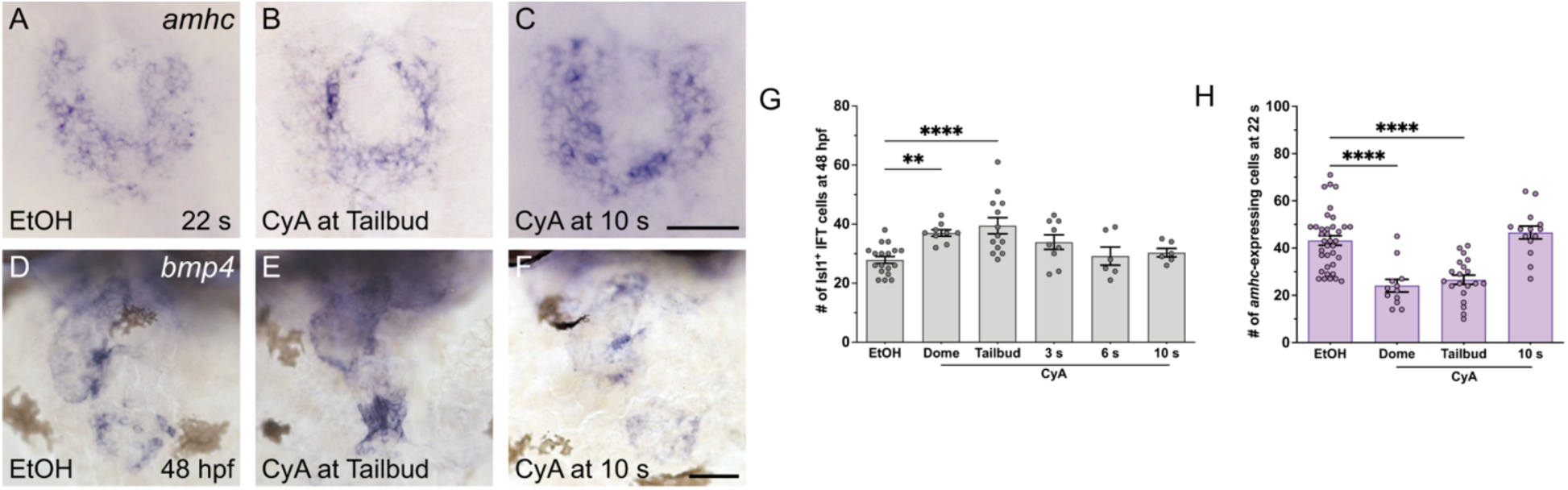
Hh signaling acts during gastrulation and early somitogenesis to limit IFT cardiomyocyte production and promote atrial cardiomyocyte production. **(A-C)** In situ hybridization depicts *amhc* expression at 22 somites (s). Dorsal views; anterior toward the top. Embryos treated with CyA at tailbud stage (B) express *amhc* in fewer cells relative to controls (A). Embryos treated with CyA at 10 s exhibit normal *amhc* expression (C). Scale bars: 50 μm. **(D-F)** In situ hybridization depicts *bmp4* at 48 hpf, frontal views. Embryos treated with CyA at tailbud (E; n=12) exhibit expanded *bmp4* expression in the IFT, relative to controls (D; n=44), whereas embryos treated at 10 s exhibit a normal pattern of *bmp4* expression (F; n=24). **(G)** Graph indicates the number of Isl1+ cardiomyocytes in the IFT at 48 hpf. Embryos treated with CyA beginning at dome or tailbud have more Isl1+ IFT cardiomyocytes, compared to ethanol-treated controls. **p<0.01; ****p<0.0001. **(H)** Graph indicates the number of *amhc*-expressing cells at 22 s. See Materials and Methods for cell counting technique. Embryos treated with CyA at dome or tailbud have fewer *amhc*-expressing cells, compared to controls. ****p<0.0001.

Interestingly, Hh signaling acts during this same early timeframe to promote production of ventricular cardiomyocytes (Thomas et al., 2008). In this previous study, we considered analysis of ventricular cardiomyocytes to be representative of both chambers, but our new IFT observations motivated us to examine whether Hh signaling also promotes atrial cardiomyocyte production during this critical time period. We treated embryos with CyA and then counted *amhc*-expressing cells at 22 s, a convenient stage for visualization of atrial cardiomyocytes prior to heart tube formation (Berdougo et al., 2003). We found that inhibition of Hh activity at dome or tailbud stage reduced the number of *amhc*-expressing cells, whereas CyA treatment at 10 s left the *amhc*-expressing population intact (Fig. 2A-C,H). These results reveal that Hh signaling plays contrasting roles during cardiac specification stages, promoting production of atrial and ventricular cardiomyocytes while simultaneously limiting the number of IFT cardiomyocytes.

### Increased Hh signaling does not affect IFT size

Our prior studies have shown that increased Hh signaling leads to excessive production of both ventricular and atrial cardiomyocytes (Thomas et al., 2008). We therefore wondered whether increased Hh signaling could also cause the loss of IFT cardiomyocytes. To examine this, we overexpressed the Hh ligand *sonic hedgehog* (*shh*) throughout the embryo via injection of *shh* mRNA (Krauss et al., 1993; Ekker et al., 1995). Consistent with our previous work (Thomas et al., 2008), *shh* overexpression increased the number of Isl1-atrial cardiomyocytes (Fig. 3A,C,F). However, the number of Isl1+ IFT cardiomyocytes did not significantly change in response to increased Hh signaling (Fig. 3A,C,F), and normal *bmp4* expression was retained in the IFT of embryos overexpressing *shh* (Fig. 3B,D). Thus, while Hh activity is necessary to prevent IFT expansion, increased Hh activity is not sufficient to depress the IFT population beyond its usual size.

**Figure 3.**
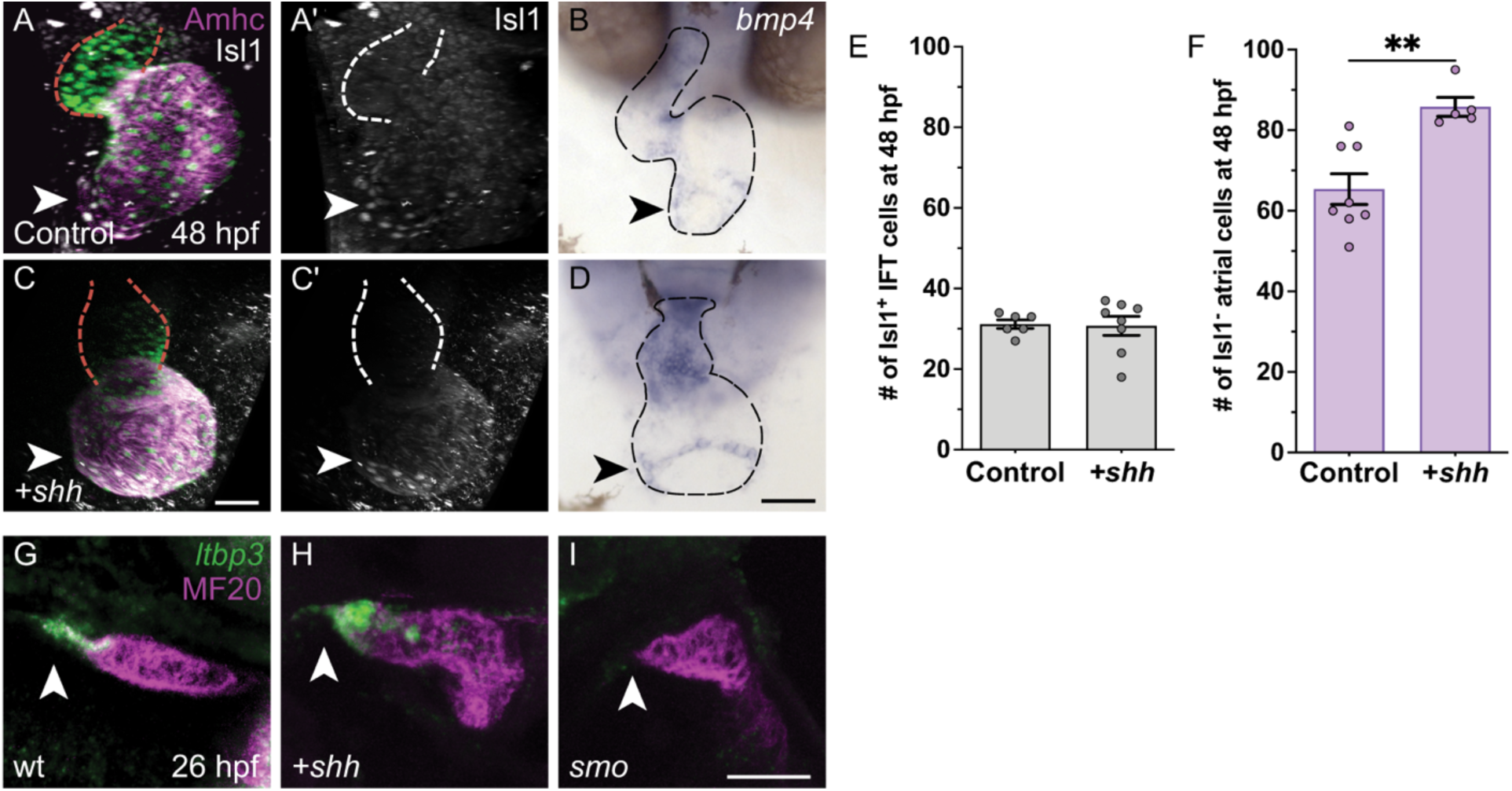
IFT cardiomyocyte number is unaffected by increased Hh signaling. **(A-D)** Control uninjected embryos (A,B) and embryos injected with *shh* mRNA (C,D) are shown after immunofluorescence (A,C) or in situ hybridization (B,D) at 48 hpf. Frontal views; arrowheads indicate the IFT. (A,C) In *Tg(myl7:H2A-mCherry)* embryos, mCherry fluorescence (green) marks cardiomyocyte nuclei, and immunofluorescence reveals Amhc (magenta) and Isl1 (white) localization; red dashed lines outline the ventricle. (A’,C’) Isl1 localization is shown in white; white dashed lines outline the ventricle. Although injection with *shh* mRNA alters cardiac morphology, the population of Isl1+ cardiomyocytes is comparable in injected embryos and controls. (B,D) Embryos injected with *shh* mRNA retain *bmp4* expression in the IFT (n=12) in a narrow ring similar to that in uninjected controls (n=9); black dashed lines outline the heart. Scale bars: 50 μm. **(E,F)** Graphs indicate the number of Isl1+ cardiomyocytes in the IFT (E) and the number of Isl1-atrial cardiomyocytes (F) at 48 hpf. See Materials and Methods for cell counting technique. Injection of *shh* mRNA increases the number of Isl1-atrial cardiomyocytes but does not change in the number of Isl1+ IFT cardiomyocytes relative to uninjected controls. **p<0.01. **(G-I)** Fluorescent in situ hybridization indicating expression of *ltbp3* (green) is combined with MF20 immunofluorescence (magenta); lateral views at 26 hpf, arrowheads indicate the arterial pole. In wt (G), *ltbp3* is expressed in OFT progenitor cells located at the arterial pole (n=13). In embryos injected with *shh* mRNA (H), *ltbp3* expression is expanded (n=9). In *smo* mutants (I)*, ltbp3* expression is typically absent (n=7/12) or reduced (n=4/12).

This finding contrasts sharply with the positive response of ventricular and atrial cardiomyocyte populations to increased Hh signaling, as mentioned above ((Thomas et al., 2008); Fig. 3A,C,F). Additionally, we found that *shh* overexpression resulted in expanded expression of *ltbp3* (Fig. 3G,H), which marks OFT progenitors in the SHF (Zhou et al., 2011). Since OFT, ventricular, and atrial populations are also reduced in *smo* embryos ((Thomas et al., 2008; Hami et al., 2011); Fig. 3I), it seems that these three cell types all respond in a reciprocal fashion to loss and gain of Hh activity. In contrast, the IFT population expands when Hh signaling is reduced but appears unaffected when Hh signaling is increased. This suggests that there are two separable roles for Hh signaling during cardiac specification stages: a dose-dependent role in promoting production of atrial, ventricular, and OFT cardiomyocytes, and a distinct permissive role that inhibits production of IFT cardiomyocytes.

### Bmp signaling acts in opposition to Hh signaling during IFT cardiomyocyte production

We wondered whether IFT cardiomyocyte production is governed by a balance between repressive and inductive signaling pathways, with the Bmp signaling pathway potentially counteracting the restrictive role of the Hh pathway. Our prior studies have demonstrated that Bmp signaling promotes production of atrial cardiomyocytes (Marques and Yelon, 2009), and additional prior work in zebrafish has shown that pharmacological inhibition of Bmp signaling can reduce the number of *isl1*-expressing cardiomyocytes at the venous pole of the atrium (Fukui et al., 2018). Moreover, our previously acquired fate map data have indicated that atrial progenitor cells reside in a ventral portion of the cardiogenic territory at 40% epiboly (Keegan et al., 2004), and retrospective analysis of these data suggests that the IFT progenitors tend to be found near the ventral edge of this territory (Fig. S1). This location places putative IFT progenitor cells in position to receive high levels of Bmp signaling during cardiac specification stages, when Bmp pathway activity is distributed in a ventral-to-dorsal gradient across the embryo (Tucker et al., 2008).

To evaluate a potential interaction between Hh and Bmp signaling during IFT formation, we utilized *lost-a-fin* (*laf*) mutants. The *laf* locus encodes the Bmp type I receptor Alk8/Acvr1l (Bauer et al., 2001; Mintzer et al., 2001); thus, in embryos homozygous for mutations in both *smo* and *laf*, both Hh and Bmp signaling would be reduced. We anticipated that classical epistasis between *smo* and *laf* would result in *smo;laf* double mutants resembling either *smo* mutants or *laf* mutants. Alternatively, parallel activities of the two pathways, such as convergence on the same target genes, could result in *smo;laf* double mutants displaying an intermediate phenotype. Indeed, in contrast to the expanded population of Isl1+ IFT cardiomyocytes observed in *smo* mutants (Fig. 4C,E,G) and the reduced population of Isl1+ IFT cardiomyocytes observed in *laf* mutants (Fig. 4C,D,G)*, smo;laf* double mutants exhibited a population of Isl1+ IFT cardiomyocytes comparable in size to their wild-type siblings at 32 hpf (Fig. 4C,F,G). Likewise, instead of the expanded or reduced expression of *bmp4* found in *smo* and *laf* mutants, respectively (Figs 1A,B,O,P and S2A,B), *smo;laf* double mutants exhibited relatively normal expression of *bmp4* in the IFT at 48 hpf (Fig. 4A,B). These results suggest that Hh signaling and Bmp signaling act in opposition to each other in order to produce an appropriately sized population of IFT cardiomyocytes.

**Figure 4.**
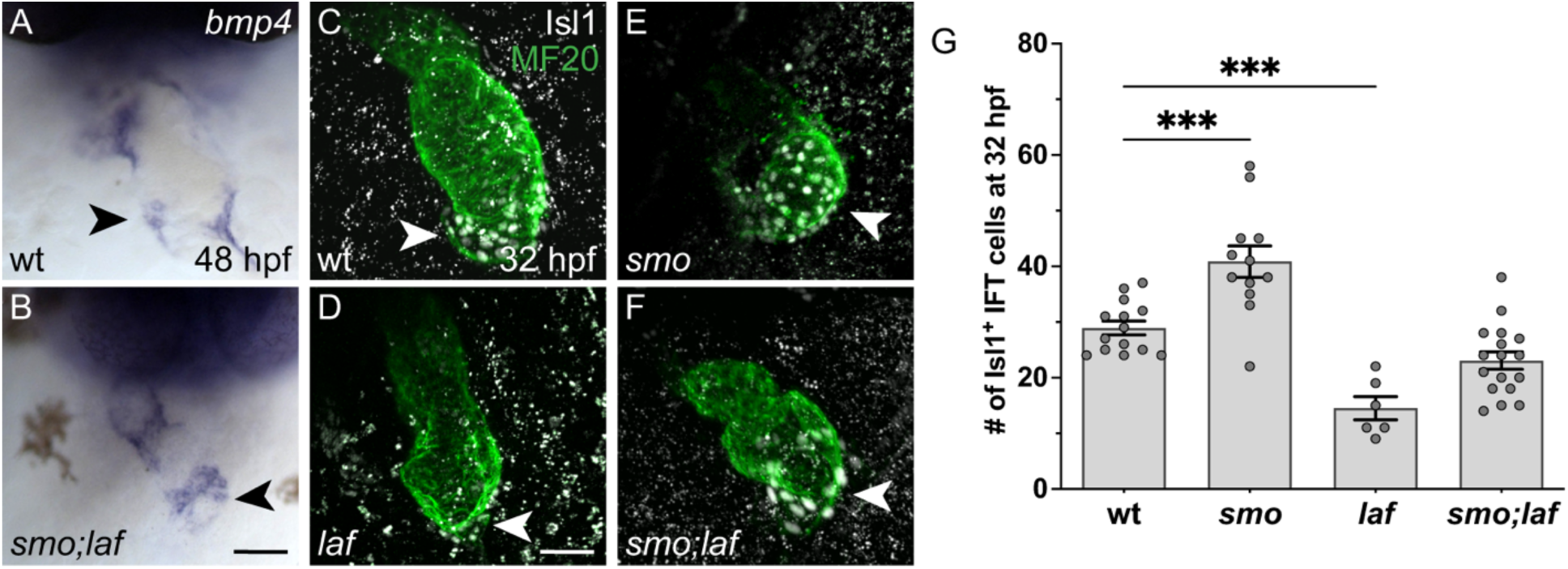
IFT cardiomyocyte number is relatively normal in *smo;laf* double mutants. **(A,B)** In situ hybridization depicts *bmp4* expression at 48 hpf. Frontal views; arrowheads indicate the IFT. *bmp4* expression is restored to a relatively normal pattern and intensity in the IFT of *smo;laf* double mutants (B, n=8), comparable to wt siblings (A, n=5). Scale bars: 50 μm. **(C-F)** Immunofluorescence depicts Isl1 (white) localization in the myocardium (MF20, green) at 32 hpf. This timepoint was chosen because *smo;laf* embryos reliably survive to 32 hpf, whereas they have variable survival at later stages. Lateral views; arrowheads indicate the IFT. Note that the arterial pole is less visible in these images than elsewhere due to the position of the heart at 32 hpf. **(G)** Graph indicates the number of Isl1+ cardiomyocytes in the IFT at 32 hpf. Isl1+ IFT cardiomyocyte number is increased in *smo* (E), decreased in *laf* (D), and relatively normal in *smo;laf* (F), relative to wt siblings (C). ***p<0.001.

### Bmp signaling acts prior to heart tube formation to promote IFT cardiomyocyte production

Next, we sought to understand the impact of Bmp signaling on IFT development in more depth. We found that *laf* mutants display a substantial reduction of Isl1+ IFT cardiomyocytes at 48 hpf (Fig. 5A-C), accompanied by diminished expression of *bmp4*, *tbx18*, and *shox2* in the IFT (Fig. S2). Additionally, *laf* mutants exhibit a small atrial chamber at this stage (Fig. 5A,B), consistent with our prior studies of *laf* (Marques and Yelon, 2009). We considered whether the decreased number of IFT cardiomyocytes in *laf* mutants might simply reflect their overall reduction in atrial size. However, this seems unlikely, since the ∼70% reduction in the number of Isl1+ IFT cardiomyocytes in *laf* mutants (Fig. 5C) was much more striking than the ∼20% decrease in the number of Amhc+ Isl1-atrial cardiomyocytes (Fig. 5D). Together with the bradycardia observed in *laf* mutants (Table S2), these data suggest an especially potent influence of Bmp signaling on IFT cardiomyocyte production.

**Figure 5.**
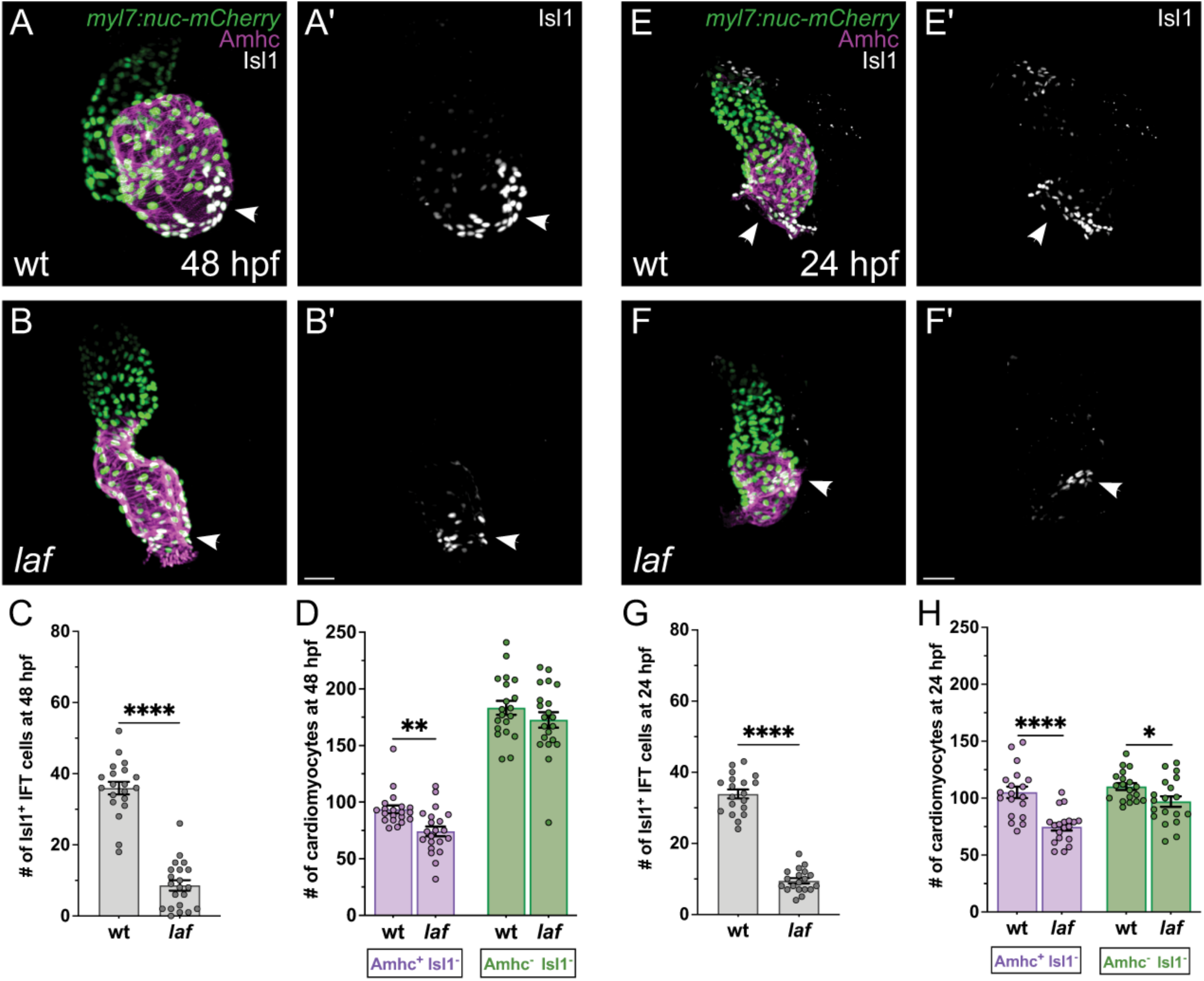
IFT cardiomyocyte number is reduced in *laf* mutants by 24 hpf (A,B,E,F) Immunofluorescence in *Tg(myl7:H2A-mCherry)* embryos depicts mCherry (green) in cardiomyocyte nuclei, as well as Amhc (magenta) and Isl1 (white) localization, in *laf* mutant embryos (B,F) and wt siblings (A,E) at 48 (A,B) or 24 (E,F) hpf. (A’,B’,E’,F’) Isl1 localization is shown in white. Frontal views, arrowheads indicate the IFT. At both stages, *laf* mutants exhibit fewer Isl1+ IFT cardiomyocytes than their wt siblings. Scale bars: 30 μm. **(C,G)** Graphs indicate the number of Isl1+ IFT cardiomyocytes in *laf* mutants and wt siblings at 48 (C) and 24 (G) hpf. The number of Isl1+ IFT cells is significantly reduced in *laf* mutants at both stages. ****p<0.0001. **(D,H)** Graphs indicate the number of Amhc+ Isl1-atrial cardiomyocytes and Amhc-Isl1-ventricular cardiomyocytes in *laf* mutants and wt siblings at 48 (D) and 24 (H) hpf. See Materials and Methods for cell counting technique. At both stages, *laf* mutants have fewer atrial cardiomyocytes than their wt siblings. *p<0.05; **p<0.01; ****p<0.0001.

Further analysis of *laf* mutants at 24 hpf revealed that their number of Isl1+ IFT cardiomyocytes is already significantly reduced in the newly formed heart tube (Fig. 5E-G). Similar to our observations at 48 hpf (Fig. 5A-D), the ∼70% reduction of the IFT cardiomyocyte population in *laf* mutants at 24 hpf (Fig. 5G) was more substantial than the ∼30% reduction in atrial cardiomyocytes seen at this stage (Fig. 5H). Thus, our findings indicate that Bmp signaling acts prior to heart tube formation to promote the establishment of IFT cardiomyocytes and that the IFT cardiomyocyte population is especially sensitive to its influence.

We also noted that *laf* mutants exhibit a reduced number of ventricular (Amhc-Isl1-) cardiomyocytes at 24 hpf (Fig. 5H). However, this ventricular deficit was not observed at 48 hpf (Fig. 5D). These results suggest that Bmp signaling may influence the timing of ventricular cardiomyocyte accumulation.

### Bmp signaling acts during gastrulation to promote IFT cardiomyocyte production

Given the impact of *laf* function on IFT cardiomyocyte number prior to heart tube formation, we asked whether Bmp signaling, like Hh signaling, influences IFT size during cardiac specification stages. To assess this, we treated embryos with DMH1, a pharmacological inhibitor of Bmp Type I receptors (Hao et al., 2010). Initiation of DMH1 treatment of wild-type embryos during gastrulation (at 40% epiboly) caused a significant reduction in the number of Isl1+ IFT cardiomyocytes (Fig. 6A,B,E), similar to the reduction seen in *laf* mutants at 48 hpf (Fig. 5A-C). In contrast, treatment with DMH1 at tailbud stage or at 24 hpf did not affect IFT size, relative to DMSO-treated controls (Fig. 6C-E). These results indicate that Bmp signaling acts during gastrulation, when cardiac progenitors are likely being specified, to promote formation of IFT cardiomyocytes.

**Figure 6.**
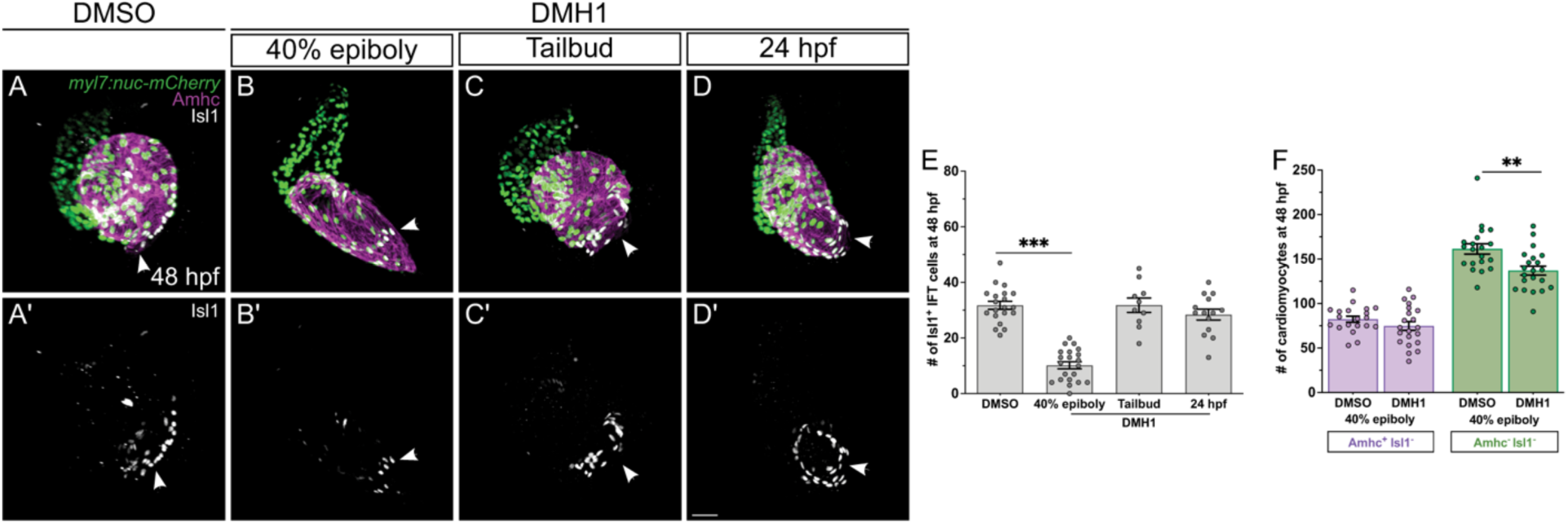
Bmp signaling acts during gastrulation to promote IFT cardiomyocyte production. **(A-D)** Immunofluorescence in *Tg(myl7:H2A-mCherry)* embryos at 48 hpf (as in Fig. 5A,B) shows effects of treatment with 1% DMSO (A), DMH1 from 40% epiboly (B), DMH1 from tailbud (C), and DMH1 from 24 hpf (D). (A’,B’,C’,D’) Isl1 localization is shown in white. Frontal views, arrowheads indicate the IFT. Treatment with DMH1 beginning at 40% epiboly reduces the number of Isl1+ IFT cardiomyocytes, reminiscent of the reduction seen in *laf* mutants (Fig. 5B,C), whereas treatment with DMH1 beginning at tailbud or 24 hpf does not seem to disrupt the formation of IFT cells. Scale bars: 30 μm. **(E)** Graph indicates the number of Isl1+ IFT cardiomyocytes in embryos treated with DMSO or with DMH1 at 40% epiboly, tailbud, or 24 hpf. ***p<0.001. **(F)** Graph indicates the numbers of Amhc+ Isl1-atrial cardiomyocytes and Amhc-Isl1-ventricular cardiomyocytes at 48 hpf in DMSO-treated controls and in embryos treated with DMH1 beginning at 40% epiboly. **p<0.01.

We note that, while DMH1 treatment at 40% epiboly yielded an IFT phenotype similar to that seen in *laf* mutants, not all aspects of the DMH1-treated hearts resembled *laf* mutant hearts. For example, DMH1-treated hearts did not display a significant reduction in Amhc+ Isl1-atrial cardiomyocytes (Fig. 6F). However, DMH1-treated embryos had significantly fewer ventricular cardiomyocytes, compared to their DMSO-treated siblings (Fig. 6F). These discrepancies may reflect differences in the specific degree and dynamics of Bmp signaling inhibition in zygotic *laf* mutants and in embryos treated with this dose of DMH1.

### Chordin acts prior to heart tube formation to restrict IFT cardiomyocyte production

Considering the loss of IFT cardiomyocyte production that is caused by inhibition of Bmp signaling, we wondered whether increased levels of Bmp signaling could expand the IFT cardiomyocyte population. To assess the impact of heightened Bmp signaling, we examined the *chordin* (*chd*) mutant phenotype. The Bmp antagonist Chordin acts by forming disulfide bonds with Bmp2/4/7, specifically interfering with their ability to bind to Bmp receptors (Piccolo et al., 1996), and zebrafish *chd* mutants exhibit increased Bmp signaling beginning in the late blastula, leading to a ventralized body pattern (Hammerschmidt et al., 1996; Schulte-Merker et al., 1997; Pomreinke et al., 2017; Zinski et al., 2017).

We found that *chd* mutants have a significant surplus of Isl1+ IFT cardiomyocytes at 48 hpf (Fig. 7A-C), and this increase is accompanied by broader expression of *bmp4* and *hcn4* at the IFT (Fig. S3). A similarly sized surplus of Isl1+ IFT cardiomyocytes is present in *chd* mutants at 24 hpf (Fig. 7E-G). This early and substantial expansion of the IFT cardiomyocyte population in *chd* mutants is reminiscent of the *smo* mutant phenotype (Fig. 1Q), and, like *smo* mutants, *chd* mutants also exhibit bradycardia (Table S3). These observations indicate that Chordin plays an important and early role in restricting IFT size and suggest that Bmp signaling promotes IFT cardiomyocyte production in a dose-dependent manner.

**Figure 7.**
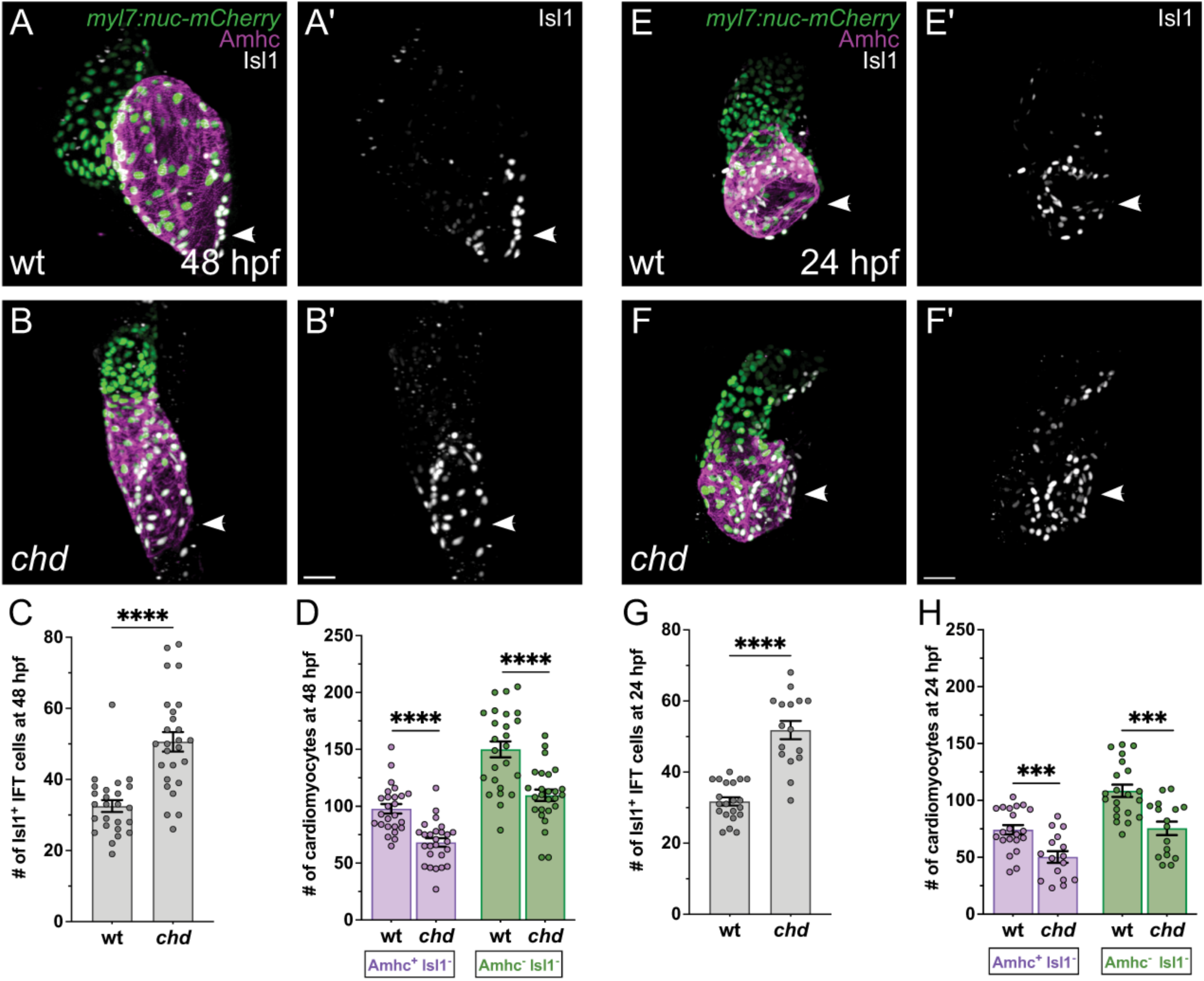
IFT cardiomyocyte number is increased in *chd* mutants by 24 hpf (A,B,E,F) Immunofluorescence in *Tg(myl7:H2A-mCherry)* embryos depicts mCherry (green) in cardiomyocyte nuclei, as well as Amhc (magenta) and Isl1 (white) localization, in *chd* mutant embryos (B,F) and wt siblings (A,E) at 48 (A,B) or 24 (E,F) hpf. (A’,B’,E’,F’) Isl1 localization is shown in white. Frontal views, arrowheads indicate the IFT. At both stages, *chd* mutants exhibit more Isl1+ IFT cardiomyocytes than their wt siblings. Scale bars: 30 μm. **(C,G)** Graphs indicate the number of Isl1+ IFT cardiomyocytes in *chd* mutants and wt siblings at 48 (C) and 24 (G) hpf. The number of Isl1+ IFT cells is significantly increased in *chd* mutants at both stages. ****p<0.0001. **(D,H)** Graphs indicate the number of Amhc+ Isl1-atrial cardiomyocytes and Amhc-Isl1-ventricular cardiomyocytes in *chd* mutants and wt siblings at 48 (D) and 24 (H) hpf. At both stages, *chd* mutants have fewer Amhc+ Isl1-atrial cardiomyocytes than their wt siblings. Additionally, *chd* mutants have fewer ventricular cardiomyocytes, compared to their wt siblings, at both stages. ***p<0.001; ****p<0.0001.

Since our prior work has indicated that heightened Bmp signaling can increase the overall size of the atrium (Marques and Yelon, 2009), we wondered whether Chordin acts to restrict the numbers of both Isl1+ IFT cardiomyocytes and Amhc+ Isl1-atrial cardiomyocytes. However, *chd* mutants have significantly fewer atrial cardiomyocytes than their wild-type siblings do at both 24 and 48 hpf (Fig. 7D,H).

Moreover, the total number of cardiomyocytes in the atrial chamber (combining the number of Isl1+ IFT cardiomyocytes and the number of Amhc+ Isl1-atrial cardiomyocytes) is relatively comparable in wild-type and *chd* mutant embryos (at 24 hpf: 106±4 in wild-type and 102±5 in *chd*; at 48 hpf: 130±4 in wild-type and 119±4 in *chd*). These results demonstrate that, although Chordin limits the size of the IFT cardiomyocyte population, it does not limit the size of the Amhc+ Isl1-atrial cardiomyocyte population. Instead, Chordin appears to act early, before the heart tube forms, to restrict the proportion of Isl1+ IFT cardiomyocytes in the atrial chamber.

We also noted a ventricular phenotype in *chd* mutants. At both 24 and 48 hpf, *chd* mutants have fewer ventricular cardiomyocytes than their wild-type siblings do (Fig. 7D,H). These observations suggest an additional role for Chordin in promoting the establishment of ventricular cardiomyocytes prior to the assembly of the heart tube.

### Bmp signaling and Hh signaling have distinct relationships with Wnt signaling during IFT cardiomyocyte production

Our data implicate Bmp signaling in supporting IFT progenitor specification (Figs 5-7), and prior studies have shown that Wnt signaling drives IFT cardiomyocyte differentiation (Ren et al., 2019). We therefore examined the relationship between the Bmp and Wnt pathways during IFT cardiomyocyte production. Hypothesizing that the early impact of Bmp signaling on IFT cardiomyocyte formation is upstream of the influence of Wnt signaling on this process, we chose to investigate whether the expansion of the IFT cardiomyocyte population observed in the context of heightened Bmp signaling is sensitive to Wnt signaling inhibition. To evaluate this, we treated embryos with PNU-7465 (PNU), an inhibitor of canonical Wnt signaling (Trosset et al., 2006).

Treatment of wild-type embryos with PNU beginning at 16 s, when IFT cardiomyocyte differentiation is underway, led to a significant reduction in the number of Isl1+ IFT cardiomyocytes (Fig. 8E,F,J), as expected based on prior studies (Ren et al., 2019). Importantly, PNU treatment also suppressed the expansion of Isl1+ IFT cardiomyocytes in *chd* mutants (Fig. 8F-H,J). Thus, the impact of heightened Bmp signaling on IFT size requires Wnt pathway activity. These results are consistent with a model in which Bmp signaling and Wnt signaling act in the same pathway to support IFT cardiomyocyte production, potentially with Bmp signaling promoting the specification of IFT progenitors that later respond to Wnt signals while differentiating into IFT cardiomyocytes.

**Figure 8.**
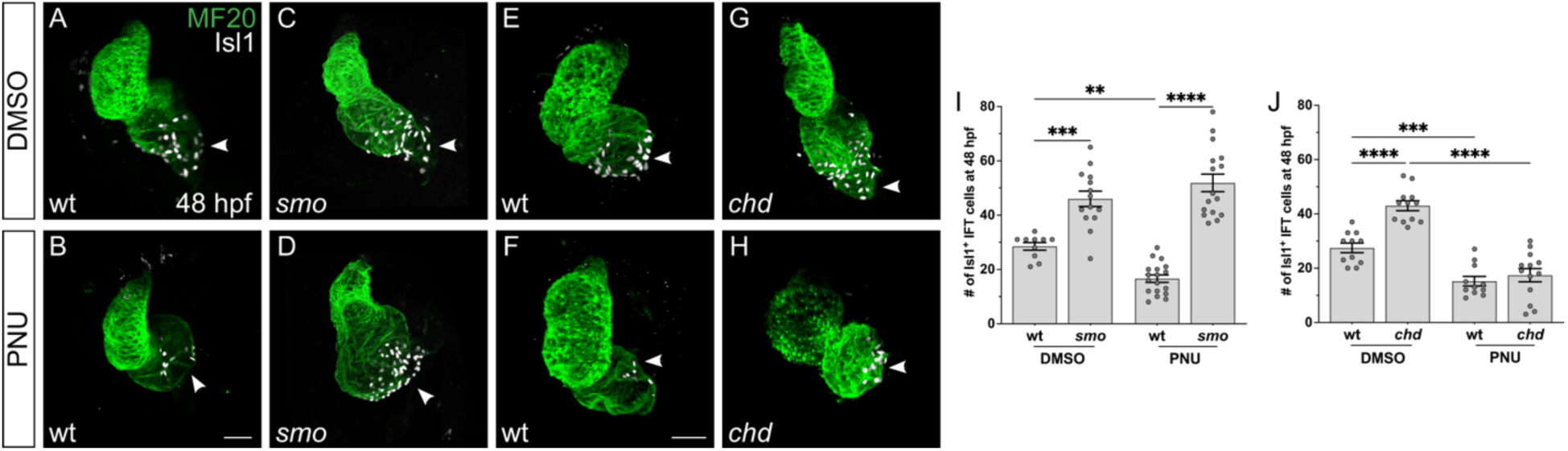
Inhibition of Wnt signaling limits IFT expansion in *chd* mutants, but not in *smo* mutants. **(A-H)** Immunofluorescence at 48 hpf depicts the effects of treatment with DMSO (A,C,E,G) or PNU-74654 (B,D,F,H) beginning at 16 s in *smo* mutants (C,D), *chd* mutants (G,H), and wt siblings (A,B,E,F). Isl1 (white) marks the IFT cardiomyocytes; MF20 marks the myocardium (green). Frontal views; arrowheads indicate the IFT. Scale bars: 30 μm. **(I,J)** Graphs indicate the effects of DMSO or PNU-74654 treatment on the number of Isl1+ IFT cardiomyocytes at 48 hpf in *smo* mutants (I) and *chd* mutants (J), with comparisons to wt siblings. Inhibition of Wnt signaling reduces the number of Isl1+ IFT cardiomyocytes in wt embryos (I,J) and in *chd* mutants (J). In contrast, *smo* mutants exhibit an increased number of Isl1+ IFT cardiomyocytes even after treatment with PNU-74654 (I). **p<0.01; ***p<0.001; ****p<0.0001.

In contrast, Wnt pathway inhibition did not alter the IFT expansion observed in *smo* mutants. PNU treatment had no significant effect on the number of Isl1+ IFT cardiomyocytes in *smo* mutants (Fig. 8C,D,I), even though PNU treatment of their wild-type siblings caused the expected reduction in IFT size (Fig. 8A,B,I). It therefore seems that Bmp signaling and Hh signaling have distinct relationships with Wnt signaling during IFT cardiomyocyte production. Our data support an early role of Hh signaling in limiting IFT progenitor specification (Figs 1,2), as well as opposing activities of Bmp and Hh signaling during IFT cardiomyocyte production (Fig. 4). However, the inability of PNU treatment to suppress the *smo* mutant phenotype suggests that Hh signaling, unlike Bmp signaling, influences IFT size through a pathway that is not dependent upon Wnt-directed IFT cardiomyocyte differentiation.

## DISCUSSION

Our studies highlight novel roles for both Hh signaling and Bmp signaling in defining the size of the cardiac IFT. Loss of Hh signaling results in an enlarged IFT, indicating that Hh signaling is required to restrict IFT cardiomyocyte number. Conversely, Bmp signaling promotes IFT cardiomyocyte production in a dose-dependent manner: reduction of Bmp signaling diminishes IFT size, while heightened Bmp signaling increases IFT size. Both of these signaling pathways exert their influence on the size of the IFT cardiomyocyte population prior to myocardial differentiation, likely during the specification of the IFT lineage. Furthermore, Hh and Bmp signaling appear to counteract each other, as inhibition of both pathways results in a relatively normal IFT. However, the interactions of these pathways with Wnt signaling during IFT formation differ: Bmp signaling functions upstream of Wnt signaling, whereas Hh signaling seems to restrict the size of the IFT through a Wnt-independent pathway. Collectively, these data support a model in which the restrictive influence of Hh signaling and the inductive influence of Bmp signaling operate in parallel and converge to specify an appropriately sized IFT progenitor pool.

This newly identified requirement for Hh signaling during IFT formation is notable for both its restrictive nature and its timing. The importance of Hh signaling for limiting the production of IFT cardiomyocytes is a striking contrast to its positive influence on the formation of other cardiomyocyte populations, including atrial and ventricular cardiomyocytes in mouse and zebrafish (Zhang et al., 2001; Thomas et al., 2008), SHF-derived OFT myocardium in mouse and zebrafish (Washington Smoak et al., 2005; Lin et al., 2006; Goddeeris et al., 2008; Hami et al., 2011), and the SHF-derived atrial septum in mouse (Goddeeris et al., 2008; Hoffmann et al., 2009; Xie et al., 2012; Briggs et al., 2016). Moreover, the restrictive influence of Hh signaling on IFT cardiomyocyte production occurs during and shortly after gastrulation, well before the onset of differentiation. We therefore favor the interpretation that Hh signaling constrains the specification of IFT progenitors, potentially through downstream pathways distinct from those that Hh signaling uses to promote specification or differentiation of other myocardial lineages. Future fate mapping studies will be needed to determine whether and how IFT progenitor specification expands when Hh signaling is inhibited. The excess IFT cardiomyocytes observed in Hh-deficient embryos may be the consequence of a fate transformation within the myocardial progenitor population, such as a reassignment of atrial progenitor cells to the IFT lineage. Alternatively, since IFT progenitors originate along the periphery of the cardiogenic territory in the early embryo (Fig. S1; (Ren et al., 2019)), the excess IFT cardiomyocytes may represent the recruitment of cells from non-myocardial lineages; perhaps Hh signaling prevents epicardial or pericardial progenitors, positioned just beyond the edge of the heart field (Ren et al., 2019; Prummel et al., 2022; Moran et al., 2024), from taking on IFT progenitor identity.

Our data also illuminate a previously underappreciated early role for Bmp signaling in promoting IFT formation. Our finding that this positive influence of Bmp signaling occurs during, but not after, gastrulation stages differs from the results of a previous study in which treatment with DMH1 at 14 hpf led to a reduced number of Isl1+ IFT cardiomyocytes (Fukui et al., 2018); this discrepancy may relate to the considerably higher concentration of DMH1 used previously (10 µM vs. 0.25 µM). Based on the impact of DMH1 during gastrulation, we propose that Bmp signaling, like Hh signaling, influences the specification of IFT progenitor cells. This timing aligns with our prior work demonstrating that Bmp signaling acts during cardiac specification stages to promote production of atrial cardiomyocytes (Marques and Yelon, 2009). However, we find that Bmp signaling exerts a stronger effect on the number of IFT cardiomyocytes than on the number of atrial cardiomyocytes, suggesting distinct functions of the Bmp pathway in specifying each progenitor pool. As with Hh signaling, future fate mapping will be necessary to determine how Bmp signaling directs myocardial fate assignments. Bmp signaling may influence the choice between IFT and atrial progenitor identities, but the shifts between IFT and atrial cardiomyocyte numbers in *laf* and *chd* mutants suggest that the fate transformations in these scenarios of reduced and heightened BMP signaling may be more complex.

It is important to note that our data do not indicate where Hh and Bmp signaling are required, relative to the IFT progenitor cells, during cardiac specification stages. One appealing model is that Bmp signaling plays a cell-autonomous role in promoting IFT progenitor specification, with receipt of particularly high levels of Bmp signaling resulting in assignment of IFT identity. This instructive role would be consistent with the small IFT cardiomyocyte population in *laf* mutants, the large IFT cardiomyocyte population in *chd* mutants, and the ventral position of putative IFT progenitors in our wild-type fate map data. It is also possible that Hh signaling plays a cell-autonomous role in repressing IFT fate assignment, such that progenitor cells receiving low levels of Hh signaling and robust levels of Bmp signaling would adopt IFT identity. Since Hh ligands are produced on the dorsal side of the embryo during gastrulation stages (Krauss et al., 1993; Ekker et al., 1995), it is feasible that the most ventral portion of the cardiogenic territory would have particularly low exposure to Hh signals. However, the idea that Hh signaling acts cell-autonomously to inhibit IFT fate specification conflicts with our observation that *shh* overexpression does not alter the number of IFT cardiomyocytes. Instead, the permissive role of Hh signaling during IFT formation is more compatible with a non-autonomous function, such as a requirement for Hh signaling in promoting the development of another tissue that, in turn, has a repressive influence on IFT progenitor specification. Future mosaic analyses, similar to our previous studies that revealed cell-autonomous roles for Hh signaling and Bmp signaling in promoting ventricular and atrial fate, respectively (Thomas et al., 2008; Marques and Yelon, 2009), will be needed to resolve where the Hh and Bmp pathways are required in the context of IFT formation.

Future work will also be important to identify the effectors that act downstream of Hh signaling and Bmp signaling to regulate IFT fate assignment. Since the Hh and Bmp pathways appear to act in opposition, it is plausible that they ultimately converge to control common downstream effectors, perhaps with Gli and Smad factors binding within the same enhancer but exerting opposite effects. Alternatively, Hh and Bmp signaling may act through different target genes, potentially within separate tissues. This latter option aligns with our observation that Hh signaling regulates IFT cardiomyocyte production independently of Wnt signaling, while Bmp signaling acts upstream of Wnt signaling. We propose that BMP signaling orchestrates IFT-specific patterns of gene expression that lay the foundation for their later receptivity to Wnt signaling and execution of IFT cardiomyocyte differentiation. In contrast, since the excess IFT cardiomyocytes that form in *smo* mutants seem impervious to inhibition of Wnt signaling, we speculate that Hh signaling acts to inhibit a different IFT specification pathway, through which the production of Isl1+ IFT cardiomyocytes bypasses, or involves to a lesser degree, a Wnt-regulated differentiation trajectory.

Altogether, our identification of novel, early, and contrasting influences of Hh and Bmp signaling on IFT cardiomyocyte production in zebrafish provides new insights into the network of pathways that regulate the initial specification of IFT progenitor cells. It will be worthwhile for future studies to examine whether the roles of the Hh and Bmp pathways in this context are conserved in mammals. Given the many critical roles played by Hh and Bmp signaling during embryonic patterning in mouse, conditional alleles that allow appropriate spatial and temporal control of these pathways will be necessary to facilitate evaluation of their impact on SAN progenitor specification. Furthermore, it will be valuable to interrogate whether modulation of Hh or Bmp signaling activity could improve the effectiveness of protocols that direct the *in vitro* differentiation of induced pluripotent stem cells into pacemaker cells (Kapoor et al., 2013; Birket et al., 2015; Protze et al., 2017; Schweizer et al., 2017). Finally, identifying Hh and Bmp signaling as early influences on the assignment of pacemaker cell identity may offer new insight into the potential developmental origins of congenital arrhythmias (van der Maarel et al., 2023).

## MATERIALS AND METHODS

### Zebrafish

We used the following zebrafish strains: *smo^b577^* (Varga et al., 2001)*, laf^sk42^*(Marques and Yelon, 2009), *chordin^tt250^* (Schulte-Merker et al., 1997), and *Tg(myl7:H2A-mCherry)^sd12^* (Schumacher et al., 2013). Embryos homozygous for *smo^b577^* were identified by their U-shaped somites (Varga et al., 2001). Embryos homozygous for *laf^sk42^* were identified by the absence of a ventral tail fin (Mintzer et al., 2001). Embryos homozygous for *chordin^tt250^* were identified by their expanded ventral tail fin (Schulte-Merker et al., 1997). Embryos carrying *Tg(myl7:H2A-mCherry)^sd12^* were identified by mCherry fluorescence (Schumacher et al., 2013). All zebrafish work followed protocols approved by the UCSD IACUC.

### In situ hybridization and immunofluorescence

In situ hybridization and immunofluorescence were performed as previously described (Zeng and Yelon, 2014). In situ probes used were: *bmp4* (ZDB-GENE-980528-2059), *tbx18* (ZDB-GENE-020529-2), *shox2* (ZDB-GENE-040426-1457), *hcn4* (ZDB-GENE-050420-360), *amhc* (*myh6*; ZDB-GENE-031112-1), and *ltbp3* (ZDB-GENE-060526-130). Primary and secondary antibodies used are listed in Table S4.

### Imaging

In situ images of wholemount samples were captured using Zeiss Axiocam cameras mounted on Zeiss Axioimager and Axiozoom microscopes, and images were processed using Zeiss Axiovision and Adobe Creative Suite software. Immunofluorescence images of wholemount samples were captured using a Leica SP5 confocal laser-scanning microscope (Figs 1, 3, and 4) and a Leica SP8 confocal laser-scanning microscope (Figs 5-8) with a 25x water objective and a slice thickness of 1 µm, and confocal stacks were analyzed using Imaris software (Bitplane) and ImageJ. All confocal images shown are 3D reconstructions, with pseudocolor employed to distinguish channels.

### Cell counting

To count Isl1+ IFT cardiomyocytes (Figs 1-8), we employed immunofluorescence with an anti-Isl1 antibody. Isl1 localizes to the nucleus of IFT cardiomyocytes, and cardiomyocytes are distinguishable from other cardiac cell types based on MF20 staining or expression of *Tg(myl7:H2A-mCherry)*. Though some pericardial Isl1 was observed, we only counted cells in which an Isl1+ nucleus was surrounded by MF20 staining or, in the case of *Tg(myl7:H2A-mCherry)* embryos, displayed nuclear mCherry. We occasionally observed embryos with poor quality anti-Isl1 staining, across all genotypes and without any discernible pattern; these samples were excluded from our analysis.

To count *amhc*-expressing cells at 22 s (Fig. 2), we used an established protocol for counting cells after in situ hybridization (Thomas et al., 2008). A cell was counted only if its nucleus was clearly outlined by *amhc* expression.

To count atrial or ventricular cardiomyocytes (Figs 3 and 5-8), we used the transgene *Tg(myl7:H2A-mCherry)* to label the nuclei of all cardiomyocytes, an anti-Isl1 antibody to label the nuclei of all IFT cardiomyocytes, and the S46 antibody (anti-Amhc) to label all atrial cardiomyocytes. To count atrial cardiomyocytes, we determined the number of mCherry+ nuclei found in Amhc+ cells and then subtracted the number of Isl1+ Amhc+ cells to attain the number of Isl1-Amhc+ cells. To count ventricular cardiomyocytes, we determined the number of mCherry+ nuclei that were in cells lacking Amhc.

### Drug treatments

Embryos were treated with CyA (Fisher Scientific 50-760-5; dissolved in ethanol) at 25-75 µM, DMH1 (Sigma-Aldrich D8946; dissolved in DMSO) at 0.25 µM, or PNU-74654 (Sigma-Aldrich P0052; dissolved in DMSO) at 25 µM in E3 embryo medium. Embryos treated with 25-75 µM CyA at dome stage (4.3 hpf) resembled *smo* mutants based on their diminutive head, mild cyclopia, ventral body curvature, and U-shaped somites. Embryos treated with 0.25 µM DMH1 at 40% epiboly (5 hpf) stage resembled *laf* mutants based on their mild dorsalization, including absence of the ventral tail fin. Embryos treated with 25 µM PNU at 16 s (17 hpf) resembled PNU-treated embryos from our previous studies (Ren et al., 2019) based on their reduced number of Isl1+ IFT cardiomyocytes (Fig. 8I,J). Control embryos were treated with an equal volume of the appropriate small molecule vehicle, either ethanol or DMSO.

### Injection of RNA

Embryos were injected with 50 pg mRNA encoding zebrafish *shha* (referred to as *shh*) at the one-cell stage (Ekker et al., 1995).

### Fate map analysis

Previously generated data sets were retrospectively analyzed to approximate the locations of IFT progenitor cells at 40% epiboly. Previously published data (Keegan et al., 2004; Keegan et al., 2005) were combined with additional unpublished data from other experiments performed in parallel. For each experimental embryo, available data indicated the initial position of the labeled cells and the subsequent location of the labeled progeny within the MF20-labeled myocardium at 48 hpf. Because IFT molecular markers were not used when generating these data sets, we used morphological criteria to score the contribution of labeled progeny to the IFT. Specifically, we scored progeny as IFT cardiomyocytes if they were located in the bottom 30% of the atrium, closer to the venous pole than to the atrioventricular canal. This approach generates a fate map with broader resolution relative to experiments in which specific molecular markers are used, but it nevertheless suggests a potential location for IFT progenitors. Instances in which the available information was insufficient to judge IFT contribution were excluded from this analysis.

### Statistics and replicates

All data shown represent at least two independent replicates from separate crosses performed on different dates. In scatterplots, each dot represents an individual sample. In bar graphs, bar height indicates the mean value for a data set, and error bars represent the standard error of the mean. All statistical analyses of data were performed using GraphPad PRISM. For each data set, a Shapiro-Wilk test was used to test normality of data. If data were normally distributed, a Student’s *t*-test (two-tailed) was performed to compare 2 sets of data, or single factor ANOVA followed by Bonferroni’s Multiple Comparison Test was performed to compare >2 sets of data. If the data were not normally distributed, a Mann-Whitney *U*-test was performed to compare data sets. Asterisks in graphs are used to indicate statistical significance: * to indicate p<0.05, ** to indicate p<0.01, *** to indicate p<0.001, and **** to indicate p<0.0001.

## ACKNOWLEDGMENTS

We thank T. Vajdi, W.E. Gordon, and M. Seifi for assistance with experiments; M. Mullins for providing *chordin* mutants, T. Sanchez, A. Yarbrough, and the UCSD Animal Care Program for zebrafish care; and members of the Chi and Yelon labs for thoughtful input.

## AUTHOR CONTRIBUTIONS

Conceptualization: R.A.R., H.G.K., D.Y.; Methodology: R.A.R., H.G.K., J.R., D.Y.; Formal analysis: R.A.R., H.G.K., C.L., D.Y.; Investigation: R.A.R., H.G.K., C.L.; Resources: J.R., N.C.C.; Writing - original draft: R.A.R., H.G.K., D.Y.; Writing - review & editing: R.A.R., H.G.K., C.L., J.R., N.C.C., D.Y.; Supervision: N.C.C., D.Y.; Project administration: D.Y.; Funding acquisition: R.A.R., H.G.K., C.L., N.C.C., D.Y.

## SUPPLEMENTARY FIGURES AND TABLES

**Supplementary Figure S1.**
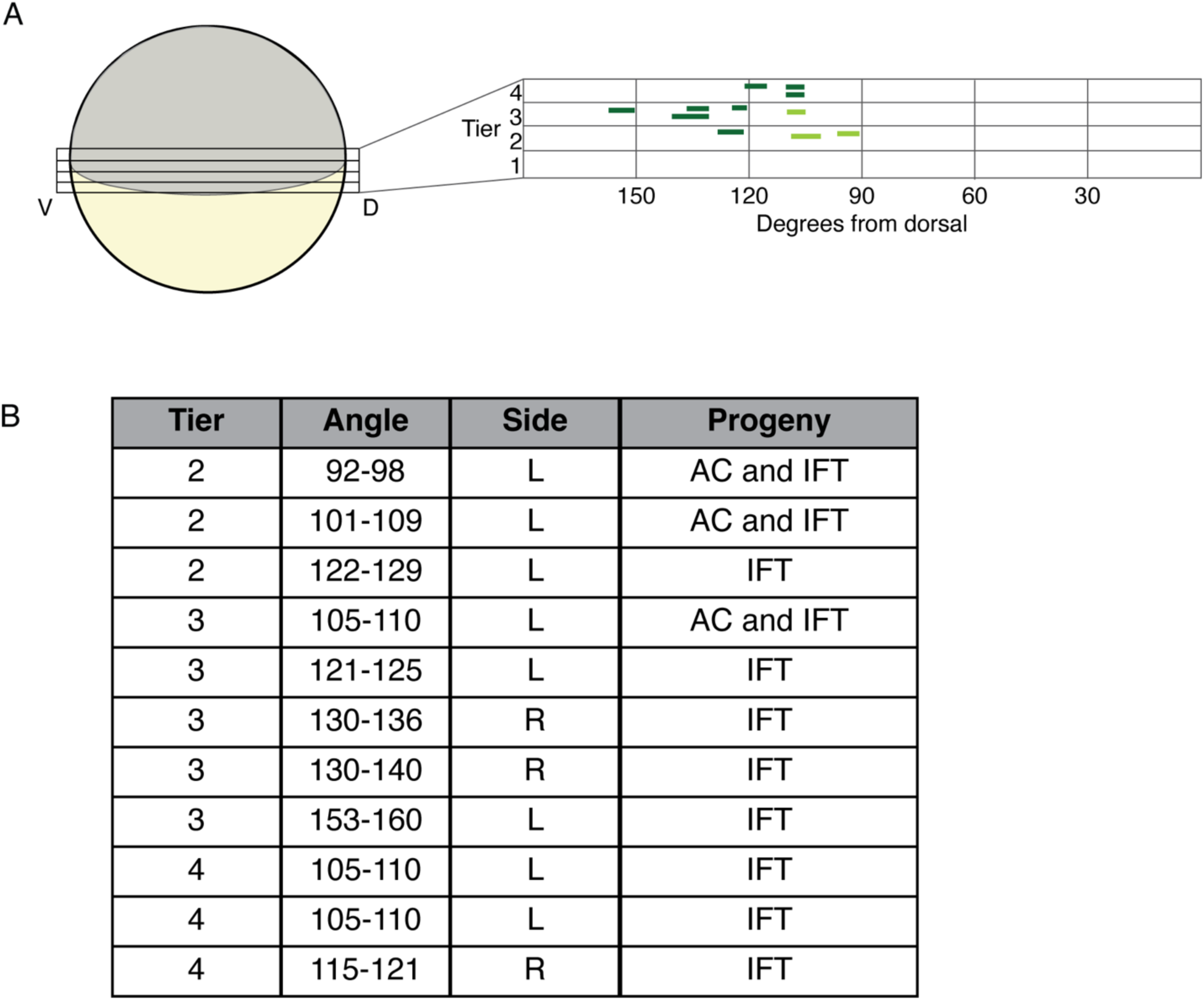
IFT progenitors originate from a ventral portion of the lateral margin. **(A)** In our previous experiments (Keegan et al., 2004; Keegan et al., 2005), fate mapping of cardiac progenitors was performed by injecting a caged fluorescein-dextran lineage tracer at the single-cell stage, followed by uncaging to label small groups of blastomeres at 40% epiboly before analysis of cell fates at 48 hpf. For each set of labeled cells, their location at 40% epiboly was recorded in terms of latitude, using cell tiers to measure distance from the margin, and longitude, using degrees from the dorsal midline. Our retrospective analysis of these data evaluated whether any labeled cardiomyocytes were found in the bottom 30% of the atrium at 48 hpf (see Materials and Methods). As this region approximates the location of Isl1+ IFT cardiomyocytes, we classified these examples as instances of IFT progeny, whereas labeled cells elsewhere in the atrium were classified as instances of atrial cardiomyocyte (AC) progeny. Altogether, we identified eight examples (marked in dark green) in which labeled cells appeared to give rise to IFT progeny. We also found an additional three examples (light green) in which labeled cells appeared to give rise to both IFT and AC progeny. In all 11 examples, IFT progenitors were located in tiers 2-4, between 92 and 160 degrees from dorsal. **(B)** Table lists each instance of IFT labeling, indicating the cell tier, degrees from dorsal, and side (left or right indicated by L or R, respectively) at 40% epiboly, as well as the final myocardial contribution of the labeled progeny at 48 hpf. We do not observe any asymmetry between the left and right sides of the embryo in this set of data: of all experiments that resulted in labeled atrial cells, we observe labeled IFT cardiomyocytes 73% (8/11) of the time on the left side of the embryo and 75% (3/4) of the time on the right side of the embryo.

**Supplementary Figure S2.**
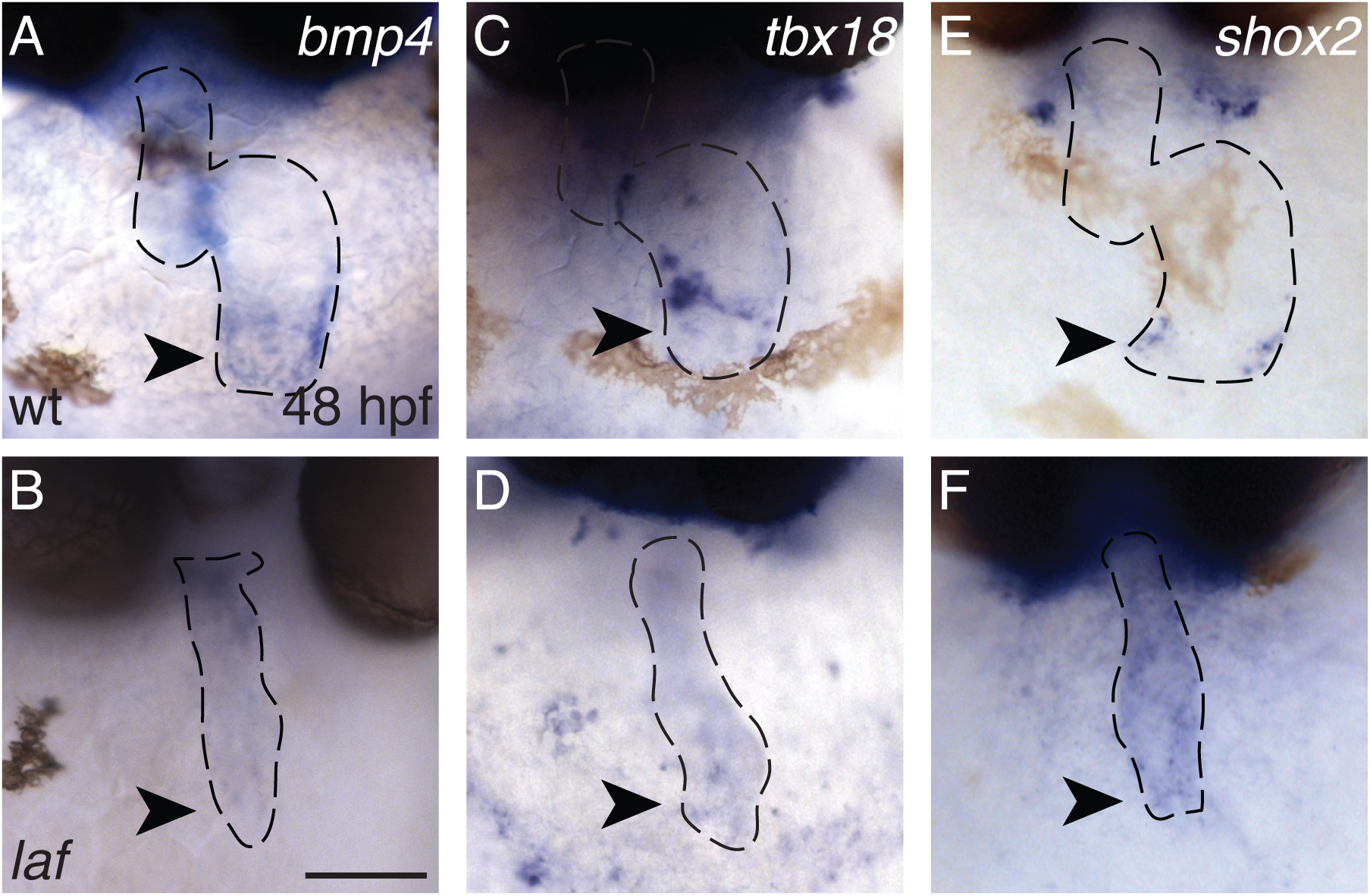
Bmp signaling promotes expression of IFT markers (A-F) In situ hybridization depicts expression of *bmp4* (A,B), *tbx18* (C,D) and *shox2* (E,F) at 48 hpf. Frontal views; arrowheads indicate the IFT. Whereas wt embryos (A,C,E) show discrete expression of IFT markers in the venous pole, concentrated expression of *bmp4* (B)*, tbx18* (D), and *shox2* (F) is not evident at the venous pole of most *laf* mutant embryos (n=44 wt, 29/52 *laf* for *bmp4;* n=11 wt, 10/11 *laf* for *tbx18;* n=13 wt, 14/14 *laf* for *shox2*). Note that variability observed in the *laf* mutant phenotype may reflect a variable degree of maternal *acvr1l* contribution in individual embryos (Mintzer et al., 2001). Scale bar: 50 μm.

**Supplementary Figure S3.**
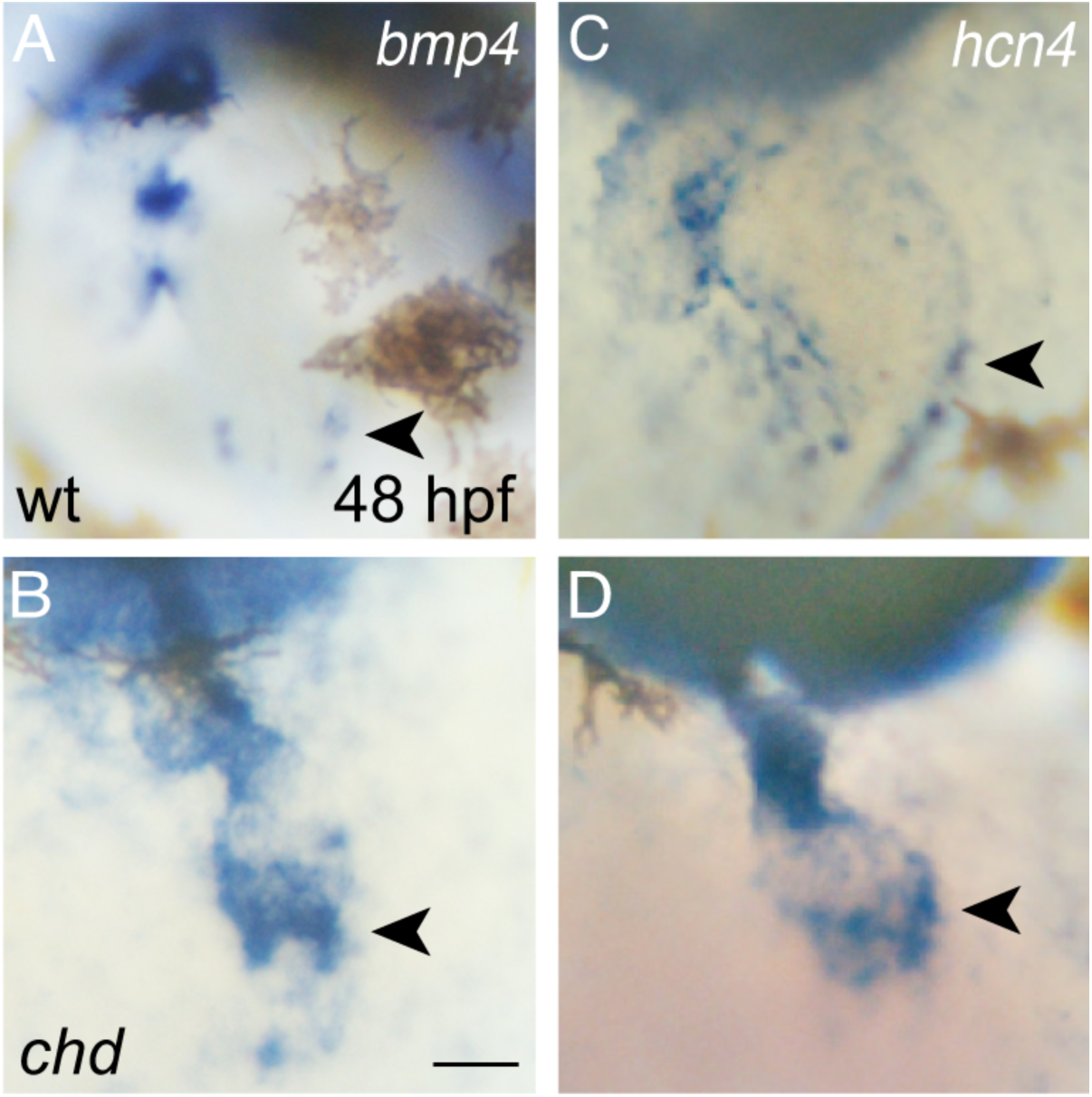
Expanded expression of IFT markers in *chd* mutants. **(A-D)** In situ hybridization depicts expression of *bmp4* (A,B) and *hcn4* (C,D) at 48 hpf in wt (A,C) and *chd* mutant embryos (B,D). Frontal views; arrowheads indicate the IFT. Broader expression of *bmp4* is visible at the IFT in *chd* mutants (n=21), compared to the expression at the IFT in wt siblings (n=24). Expression of *hcn4* expression is similarly expanded in *chd* mutants (n=16), compared to wt siblings (n=19). Scale bar: 30 μm.

**Supplementary Table S1.** Slow heart rate in *smo* mutants

| Genotype | Heart rate (bpm) | n |
| --- | --- | --- |
| wild-type | 116 ± 1 | 42 |
| *smo* mutant | 79 ± 2 | 41 |

*smo* mutants exhibited a significantly reduced average heart rate (beats per minute (bpm) ± standard error) compared to their wild-type siblings ( $p < 0.0001$ ) at 48 hpf. Additionally, a subset of *smo* mutants (3/41) exhibited arrhythmia.

**Supplementary Table S2.** Slow heart rate in *laf* mutants

| Genotype | Heart rate (bpm) | n |
| --- | --- | --- |
| wild-type | 100 ± 2 | 27 |
| *laf* mutant | 62 ± 4 | 27 |

*laf* mutants exhibited a significantly reduced average heart rate (± standard error) compared to their wild-type siblings ( $p < 0.0001$ ) at 36 hpf.

**Supplementary Table S3.** Slow heart rate in *chd* mutants

| Genotype | Heart rate (bpm) | n |
| --- | --- | --- |
| wild-type | 141 ± 2 | 16 |
| *chd* mutant | 118 ± 7 | 14 |

*chd* mutants exhibited a significantly reduced average heart rate (± standard error) compared to their wild-type siblings ( $p = 0.0012$ ) at 48 hpf.

**Supplementary Table S4.** Antibodies used for immunofluorescence

| **Name** | **Source** | **Catalog #** | **Dilution** |
| --- | --- | --- | --- |
| **Primary antibodies** |  |  |  |
| MF20 hybridoma supernatant | DSHB | MF20 | 1:10 |
| S46 hybridoma supernatant | DSHB | S46 | 1:10 |
| Rabbit anti-Isl1 polyclonal | GeneTex | 128201 | 1:2000 |
| Rat anti-mCherry mAb, Alexa Fluor 594 | ThermoFisher Scientific | M11240 | 1:400 |
| **Secondary antibodies** |  |  |  |
| Goat anti-Mouse IgG2b, TRITC | Southern Biotech | 1090-03 | 1:100 |
| Goat anti-Mouse IgG1, FITC | Southern Biotech | 1070-02 | 1:100 |
| Goat anti-Rabbit IgG (H+L), Alexa 488 | ThermoFisher Scientific | A11008 | 1:500 |
| Goat anti-Rabbit IgG (H+L), Alexa 647 | ThermoFisher Scientific | A21245 | 1:500 |
| Goat anti-Mouse IgG (H+L), Alexa 647 | ThermoFisher Scientific | A21235 | 1:100 |

